# Chill tolerant *Drosophila* species maintain electrogenic muscle membrane potential to resist cold-induced depolarization

**DOI:** 10.1101/2025.02.26.640316

**Authors:** Johannes Overgaard, Jeppe Seamus Bayley, Jacob Nørgaard Poulsen, Nikolaj Johannes Skole Jensen, Thomas Holm Pedersen, Jon Herskind, Mads Kuhlmann Andersen

## Abstract

The ability to tolerate low temperature is among the most important traits defining the functional niche of insects and it clear that cold tolerance of most insects is intimately linked to their ability to defend membrane potential (V_m_). Failure to maintain membrane polarization results in loss of neuromuscular function and may ultimately initiate cell death and organismal injury. Prolonged cold exposure challenges membrane polarization through loss of transmembrane ion balance; however, the insect muscle V_m_ is also dependent on a strong and temperature-dependent electrogenic effect driven by Na^+^/K^+^-ATPase activity. In the present study we investigate the electrogenic contribution of the Na^+^/K^+^-ATPase at benign (20°C) and low (0°C) temperature in ten *Drosophila* species representing a broad spectrum of chill tolerance. We find that the electrogenic effect of the Na^+^/K^+^-ATPase contributes a considerable component of the muscle V_m_ in all ten species at 20°C. This electrogenic contribution is reduced significantly at 0°C in the chill sensitive species, while tolerant species retain their electrogenic effect at low temperature. Thus, the initial cold-induced muscle depolarization, that is a hallmark of chill sensitive insects, is largely caused by loss of Na^+^/K^+^-ATPase-dependent electrogenic polarization. We hypothesized that maintenance of Na^+^/K^+^-ATPase activity in the cold would be energetically costly, but in contrast to our hypothesis we find no evidence for major energetic costs in the species that maintain membrane polarization at low temperature. On the basis of these observations we discuss how other adaptations at the protein or membrane level could explain the observed intraspecific differences.

## Introduction

When subjected to stressful cold, chill susceptible insects are challenged on their ability to maintain physiological homeostasis including maintenance of ion balance, membrane potential, excitability and cellular viability (Koštál et al., 2004; MacMillan and Sinclair, 2011a; Overgaard and MacMillan, 2017; Overgaard et al., 2021). The categorization of “chill susceptible” insects includes the majority of insect species and covers both species that are stressed at mild temperature (chill sensitive) and those that defend homeostasis and function well below 0°C (chill tolerant) (Lee, 1991; Bale, 1996; Sinclair, 1999). Even though the critical temperature to disturb physiological homeostasis varies considerably among species, most studies suggest that the physiological syndrome of cold stress is the same once the species-specific critical temperature is reached (Koštál et al., 2007; MacMillan et al., 2015a; Des Marteaux and Sinclair, 2016; Andersen et al., 2017b).

Exposure to stressful cold in chill susceptible insects is characterized by an archetypical loss of organismal performance that is tightly linked to the loss of nervous and muscular membrane polarization and excitability (Overgaard and MacMillan, 2017; Overgaard et al., 2021). Initially cold exposure triggers the onset of a chill coma caused by spreading depolarization in the central nervous system (Robertson et al., 2017; Andersen et al., 2018; Robertson et al., 2023). This chill coma is reversible if the insects are returned to permissive temperatures. However, during longer and/or more intense cold exposures the organismal homeostasis is further disrupted. First, exposure to low temperature in itself causes a depolarization of the muscle membrane potential (V_m_) which compromises muscle function and exacerbates the coma (Goller and Esch, 1990; Hosler et al., 2000; MacMillan et al., 2014; Andersen et al., 2015; Findsen et al., 2016). Secondly, prolonged cold exposure impairs epithelial and osmoregulatory function causing a gradual disruption of extracellular ion homeostasis characterized by hemolymph hyperkalemia (MacMillan and Sinclair, 2011b; Overgaard and MacMillan, 2017; MacMillan, 2019). This hyperkalemia causes further membrane depolarization and ultimately the combined effects of cold and hyperkalemia can initiate cell death, likely through opening of voltage gated Ca^2+^ channels (MacMillan et al., 2015b; Bayley et al., 2018; Carrington et al., 2020). Accordingly, it is clear that the ability to mitigate the loss of homeostasis and maintain well-polarized excitable tissues at low temperature is defining for the chill tolerance of chill susceptible insects. Indeed, numerous studies have demonstrated how adaptation (across generations) and/or acclimation (within an individual) effects on muscle-, neuro- and/or osmoregulatory act to preserve cellular and extracellular homeostasis are closely linked to the level of organismal chill tolerance among insects (reviewed in Overgaard and MacMillan (2017) and Overgaard et al. (2021)).

The role of low temperature on cellular membrane potential (V_m_) in insects is influenced by active and passive membrane properties components that, for simplicity, can be partitioned into a diffusional component and an electrogenic component (Overgaard and MacMillan, 2017), but in reality these components are integrated and co-dependent as explained by the ‘charge difference’ model (Fraser and Huang, 2004; Bayley et al., 2021). The diffusional component includes all the variables in the Goldman-Hodgkin-Katz equation that describes the cell’s resting membrane potential (Goldman, 1943; Hodgkin and Katz, 1949; Hodgkin and Horowicz, 1959). Thus, cold exposure reduces the diffusional component of membrane potential directly and thermodynamically through the change in absolute temperature, but also through the large disruptions in ion balance characterized principally by the cold-induced hyperkalemia (Overgaard and MacMillan, 2017; MacMillan, 2019). Interestingly, very little is known of how cold affects relative permeability which could potentially also alter how insect membrane polarization is affected by cold.

The other main component of cell polarization in insect muscle is linked to the electrogenic effect generated by the active transport of ions across the muscle membrane (Thomas, 1972; Skou and Esmann, 1992). Most notably, the Na^+^/K^+^-ATPase generates a net current from the uneven movement of cations across the membrane (i.e. 3 Na^+^ out and 2 K^+^ in). This produces an electrogenic potential and according to the insights from Ohm’s law it follows that this electrogenic current produces a potential that is proportional to the magnitude of both the current and the membrane resistance (Glitsch, 2001). The magnitude of the electrogenic component differs between species, but at room temperature this component has been reported to contribute considerably (c. 10 to 20 mV) to the resting muscle membrane potential in insects, *Drosophila* included (Rheuben, 1972; Wareham et al., 1974; Henon and Ikeda, 1981; Bayley et al., 2020). It is also clear that acute lowering of temperature will reduce the magnitude of active ion transport and thus cause a rapid depolarization of muscle V_m_. Indeed, previous measurements of insect muscle V_m_ have demonstrated a strong depolarizing effect with acute cooling (Wareham et al., 1974; MacMillan et al., 2014; Andersen et al., 2017b; Bayley et al., 2020), and this is also true for *Drosophila* (Andersen et al., 2015; Cheslock et al., 2021). Furthermore, the study by Cheslock et al. (2021) revealed that cold acclimation alleviated the thermal dependency of the electrogenic effect, possibly due to changes in Na^+^/K^+^-ATPase activity.

By comparing *Drosophila* species, ranging from temperate to tropical, we have previously found large interspecific differences in the ability to maintain muscle V_m_ during cooling (Andersen et al., 2015; Andersen and Overgaard, 2019). Thus, chill tolerant *Drosophila* species maintain cellular polarization down to 0°C, while chill sensitive species lose cellular polarization around 10°C and experience a marked depolarization at 0°C. However, our previous measurements were performed *in vivo*, without control of the extracellular environment, and the differential roles of diffusion potential (including hyperkalemia) and the electrogenic effects therefore remain unresolved. The aim of the present study was to investigate if chill tolerant species are better able to preserve membrane polarization at low temperatures, and specifically to examine if preservation of the electrogenic component contributes to such cold adaptation. Further, because the electrogenic polarization is fueled by active membrane transport we examined if species defending membrane polarization at low temperature are characterized by a relative higher energy metabolism during cold exposure. To specifically address these questions, we quantified the Na^+^/K^+^-ATPase-dependent electrogenic effect of muscle V_m_ at high (20°C) and low (0°C) temperature in ten *Drosophila* species by applying the specific blocker ouabain. The ten species used differ markedly in their chill tolerance and some are known to maintain cell polarization at low temperature *in vivo* (Andersen et al., 2015; Andersen and Overgaard, 2019). Further, we examined if chill tolerant species are characterized by a relatively higher aerobic metabolism at low temperature which could putatively be linked to sustained Na^+^/K^+^-ATPase activity under these conditions.

## Materials and methods

### Animal husbandry

The ten *Drosophila* species used in this study were chosen to represent a gradient of chill tolerance based on Kellermann et al. (2012) (see **Table 1**). Parental populations of the ten species were maintained under common garden conditions at a constant 19 °C and a long day light regime (18/6 h light/dark cycle while flies were sampled for measurement of membrane potentials, which was changed to 22/2 h light/dark for unrelated reasons before metabolic rate measurements). Flies were kept in bottles with 40 mL oatmeal-based Leeds medium (ingredients per liter water: 60 g yeast, 40 g sucrose, 30 g oatmeal, 16 g agar, 12 g methylparaben, and 1.2 mL acetic acid). Depending on the species, adult flies were allowed to oviposit for 2-4 days on fresh medium in a new bottle to ensure appropriate rearing densities of the offspring (MacMillan et al., 2015a). Emerging flies were moved to new bottles and left to mature for seven days before experiments (at the age of 7-8 days). Due to their larger size, only female flies were used in these experiments.

**Table 1.** Descriptions and chill coma temperatures (CT_min_) for *Drosophila* species used in this study. Data from Kellermann et al. (2012).

| Species | Subgenus | Group | Subgroup | CT<sub>min</sub> (°C) |
| --- | --- | --- | --- | --- |
| *Drosophila montana* | *Drosophila* | *Virillis* | - | -3.13 |
| *Drosophila ezoana* | *Drosophila* | *Virillis* | - | -2.39 |
| *Drosophila littoralis* | *Drosophila* | *Virillis* | - | -1.98 |
| *Drosophila obscura* | *Sophophora* | *Obscura* | *Obscura* | -0.57 |
| *Drosophila virilis* | *Drosophila* | *Virillis* | - | 0.80 |
| *Drosophila melanogaster* | *Sophophora* | *Melanogaster* | *Melanogaster* | 2.21 |
| *Drosophila rufa* | *Sophophora* | *Melanogaster* | *Montinum* | 3.63 |
| *Drosophila teissieri* | *Sophophora* | *Melanogaster* | *Melanogaster* | 6.67 |
| *Drosophila bunnanda* | *Sophophora* | *Melanogaster* | *Montinum* | 7.34 |
| *Drosophila bipectinata* | *Sophophora* | *Melanogaster* | *Ananassae* | 7.37 |

### Experimental saline

Measurements of muscle membrane potentials *in vitro* required the use of saline, for which we modified a previously established *Drosophila* saline (Andersen et al., 2017c) to more closely reflect the hemolymph concentrations of Na^+^, K^+^, and Cl^−^ (van der Meer and Jaffe, 1983; MacMillan et al., 2015a) and osmolality (Olsson et al., 2016). Thus, the final saline contained (in mmol L^−1^): 80 NaCl, 15 KCl, 4.3 NaH_2_PO_4_, 10.2 NaHCO_3_, 2 CaCl_2_, 8.5 MgCl_2_, 10 glutamine, 15 HEPES, and 100 glucose). Saline pH was adjusted to 7.0 by titration with 4 mol L^−1^ NaOH.

### Effects of temperature on the muscle membrane potential

For measurements of membrane potentials, flies were briefly submerged in 70 % ethanol to dewax the cuticle. From here, the head was cut longitudinally to disrupt the CNS, after which the flies were transferred to a glass petri dish containing a silicone elastomer base (Sylgaard, Dow Corning Corp., Midland, MI, USA) and fixed with microneedles through the head and abdomen. Subsequently, flies were submerged in experimental saline and the dorsal longitudinal flight muscle fibers were exposed by cutting off a small part of the dorsal cuticle of the thorax using a pair of micro-scissors. The petri dish with the fly preparation was placed in a jacketed glass chamber where temperature was controlled by circulating a water-glycol mixture with a cooling bath. In all experiments, fly preparations were incubated in standard *Drosophila* saline at the desired experimental temperature (0 or 20°C) for 15-20 minutes before membrane potentials were recorded.

Membrane potentials were measured using glass microelectrodes (1B100F-4, World Precision Instruments (WPI), Sarasota, FL, USA) which were pulled in a Flaming-Brown P-97 electrode puller (Sutter Instruments, Novato, CA, USA) to a tip resistance of 10-15 MΩ when filled with 3 mol L^−1^ KCl. The microelectrode was connected to an electrode head stage in a micromanipulator and connected to a differential electrometer (Electro 705, WPI) while a chlorinated silver wire reference electrode was placed in the experimental saline away from the fly. Raw data from the electrometer was digitized with a Micro1401-3 data-acquisition system (Cambridge Electronic Design, Cambridge, UK) and collected by a computer running Spike2 software (v8, Cambridge Electronic Design, Cambridge, UK). From here, the glass microelectrode was placed next to the muscle, the potential zeroed, and the electrode advanced slowly until a sharp drop in measured potential was observed (indicative of having penetrated the muscle cell membrane), after which the electrode was retracted to return to zero. The observed drop in potential was recorded as the muscle membrane potential (V_m_) of the muscle cell if this potential remained stable over three consecutive penetrations. When possible, multiple muscle fibers were measured in each fly (1-4 measurements per fly). For most flies more than three were obtained for each experimental condition, and their average potential was noted as the muscle V_m_ for that replicate. This resulted in a total of 151 measurements of muscle V_m_ with 4-10 independent measurements for each species/temperature combination (see **Table S1** for specific sample sizes).

### Measurements of the electrogenic component

To determine the electrogenic component of the muscle V_m_, we followed the procedure outlined above, however, in these experiments, 1 mmol L^−1^ ouabain (a specific blocker of the Na^+^/K^+^-ATPase) was added to the experimental saline. Again, preparations were incubated for 15-20 minutes before measurements began. This experiment results in an additional 153 muscle V_m_ measurements in the presence of ouabain with 5-10 independent measurements for each species/temperature combination (see **Table S1**). The species- and temperature-specific magnitude of the electrogenic component (in mV) was then estimated by subtracting average muscle V_m_ recorded for a species/temperature treatment in the standard saline from the average of that measured in the presence of ouabain.

### Stop-flow respirometry

To investigate if the ability to maintain muscle membrane polarization at low temperature was correlated to increased metabolic costs at the whole organism level, we used indirect calorimetry to assess metabolic rate at eight temperatures ranging from 0 to 19°C (19, 14, 11, 9, 7, 5, 3, and 0°C). CO_2_ production rate (V[CO_2_) was measured using stop-flow respirometry in a setup as described by Jensen et al. (2014). Here, groups of female flies of each species were randomly allocated to one of 16 cylindrical glass chambers (metabolic chambers, length = 6 cm, diameter = 2 cm) and provided with a moistened piece of paper (1.5 x 3.5 cm) with 0.1mL of Leeds medium as a source for food and water. More individuals of small species (up to 92) were used per chamber compared to large species (down to 24) to ensure high quality, detectable levels of CO_2_-production at all temperatures. Two parallel 8-channels-multiplexers (RM Gas Flow Multiplexer, Sable Systems, Las Vegas, NV, USA) controlled the repeating sequential flushing (opening) of the 16 chambers such that repeated measurements of V[CO_2_ were obtained in each chamber. In each run (i.e. running the 16 chambers through all temperatures), 12 of the chambers contained flies while the remaining four were empty to measure the magnitude of background respiration/CO_2_ leakage. Airflow was secured by a vacuum pump and maintained at 150 mL min^−1^ by an adjustable mass flow meter (Side-Trak, Sierra Instruments, Monterey, CA, USA) controlled by a flow controller (MFC 2-channel v. 1.0, Sable Systems, Las Vegas, NV, USA). Before entering the metabolic chambers, atmospheric air passed a soda lime column (MERCK Millipore, Darmstadt, Germany) followed by a dried CaCl_2_-column (AppliChem, Darmstadt, Germany) to remove CO_2_ and water vapor, respectively. Air leaving the chamber passed another CaCl_2_-column to remove respiratory water vapor before entering a CO_2_-analyzer (Li7000 CO_2_/H_2_O Gas Analyzer, Li-COR Environmental, Lincoln, NE, USA). Each chamber was flushed for 2.5 min at a time, and because of the sequential flushing of individual chambers, the closing period for each chamber was 37.5 min (remaining 15 chambers x 2.5 min). At low temperature (0, 3 and 5°C), flushing and closed periods lasted longer (5 and 75 min, respectively) to ensure a good signal-to-noise ratio of CO_2_ production despite the reduced metabolic rates. To control temperature, the multiplexers and respiration chambers were placed inside a programmable temperature cabinet (KB8182, Termaks AS, Kungsbacka, Sweden) where flies were exposed to stepwise decreasing temperatures (i.e. 19, 14, 11, 9, 7, 5, 3, and 0°C). Further, to ensure that the temperature of air used for flushing chambers was equilibrated to the incubator, it was run through a copper tube coil before reaching the metabolic chambers. Flies were held at each temperature for 240 or 480 min (i.e. six sequence cycles of 40 or 80 min) at high and low temperature respectively.

During the experiments we observed that the least chill tolerant species (*D. melanogaster*, *D. rufa*, *D. teisseiri*, *D. bunnanda*, and *D. bipectinata*) became comatose at the lowest temperatures, we therefore ensured that all flies were alive (i.e. by allowing comatose flies to recover at room temperature) before the flies, were frozen at -20°C for two hours after which total fresh mass and the precise number of flies was determined. In a follow-up experiment, we determined the temperature of chill coma onset of the above five species in which paralysis was observed. Groups of flies, separated by species only, were enclosed in the metabolic chambers as described above but with free access to air supply. Subsequently, we ran the temperature protocol and at each experimental temperature we recorded if different species were able to move by gently tapping the metabolic chamber.

### Respiration data analysis

To obtain V□CO_2_ (in µL/h), CO_2_-respiration curves were integrated automatically using a script in Mathematica (version 7.0, Wolfram Research, Champaign, Illinois, USA) (see Jensen et al. (2014)). To correct for background respiration/diffusion, we deducted the miniscule assessments of V□CO_2_ we could record from the empty chambers. Mass-specific V□CO_2_ (µL/h/g) was calculated by dividing the V□CO_2_ from each metabolic chamber with the total fresh mass of flies in said metabolic chamber. Due to the changing temperature during the transition between experimental temperatures, the first two sequence cycles at each temperature was excluded to avoid measurements from time points where flies and experimental system were still adjusting to the new temperature. For the initial temperature (19°C), we removed four cycles as this allowed the flies longer time to adjust to the metabolic chambers.

V□CO_2_ was expressed relative to body mass and to account for allometric scaling we scaled the V□CO_2_ of each measurement to a fly body mass of 1 mg using the following formula:

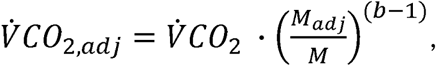

where V□*CO_2, adj_* is the scaling-adjusted, mass-specific rate of CO_2_ production, V□*CO_2_* is the original, unadjusted mass-specific V□CO_2_, *M_adj_* is the mass being adjusted to (here 1 mg), *M* is the mass of the average fly in the chamber (obtained by dividing the fresh mass of flies from each metabolic chamber with the number of flies in the respective chamber), and *b* is the allometric scaling exponent. For this analysis we used a scaling exponent *b* = 0.75 which is close to previous assessments of the allometric scaling factor in drosophilid flies (Oikawa et al., 2006; Messamah et al., 2017). In total this approach resulted in 516 independent estimates of V□CO_2_ with sample size of 4-12 for each species/temperature combination (see **Table S2** for sample sizes).

### Statistical analyses

All statistical analyses were performed in R software (v 4.2.3, R Core Team (2023)). The dataset on muscle V_m_ was tested for data normality using Shapiro Wilk’s tests. Groups that violated normality were tested for outliers using Grubb’s test (grubbs.test() function from the ‘*outliers*’ package (Komsta, 2022)), and in three of four cases where data was non-normal, removal of a single outlier resulted in normal-distributed data. The last case (*D. ezoana* at 20°C without ouabain) was used in parametric analyses as is, despite remaining non-normal distributed (i.e. one of 40 groups was non-normal). Subsequently, the effects of species, temperature, ouabain, and their interactions on muscle V_m_ were analyzed using a three-way ANOVA, which included all three parameters as fixed factors. This approach resulted in a significant three-way interaction and we therefore based our statistical analysis of this dataset on 1) dividing the dataset into two (control and ouabain treatment) and 2) the calculated electrogenic effects (i.e. the difference in V_m_ between control and ouabain-treated preparations). For each of these simplified datasets, we used linear models to test for the effects of experimental temperature and species-specific chill tolerance (CT_min_ for each species). Finally, we reduced this dataset even further by calculating the thermal dependency of the electrogenic effect (in mV per °C) and correlated this to the CT_min_ using a linear regression.

For the respirometry data, the adjusted mass-specific V[CO_2_ was first tested for data normality in the same way as the muscle V_m_ dataset. Four of 80 groups (of species x temperature) were found to violate normality. However, due to the small sample sizes for some of these groups (N = 4, 6, 6, and 12) we decided to proceed with parametric analyses. Here we tested the effects of species and temperature on the log_10_-transformed mass-specific V[CO_2_ with a linear mixed-effect model using the lme() function from the ‘*nlme*’ package. This model included species (categorical) and temperature (continuous) as fixed factors, and metabolic chamber and experimental run as random factors, with metabolic chamber nested in experimental run as each chamber was measured at multiple temperatures within the same run. Based on this dataset, we then calculated thermal quotient, Q_10_, for each species using the Q10() function from the ‘*respirometry*’ package (Birk, 2018) and did so over three temperature ranges: 1) The full temperature spectrum, 2) at low temperatures where some species enter coma and others do not (≤ 9°C), and 3) at moderate temperatures where all species are able to be active (≥ 9°C). Again, these were correlated to species-specific CT_min_ data using linear regression.

Ideally, the correlations mentioned above should account for relatedness in the *Drosophila* phylogeny. However, species were chosen not only based on their chill tolerance and phylogeny, but also considering the stability of the preparation (more species were attempted; however, their small size, and brittle cuticle complicated dissections in relation to V_m_ measurements). For example, our dataset contains several species from the same group (i.e. *Virillis*) and subgroup (see **Table 1**), resulting in a phylogenetically unbalanced design. It is difficult to address this issue considering that 10 species is already at the lower limit for appropriate phylogenetic analyses. Our solution to this dilemma was to use a non-phylogenetic approach to our correlations, but discuss this caveat in our interpretation of the data where appropriate. P < 0.05 was considered statistically significant in all analyses.

## Results

### Muscle membrane potential

Muscle membrane potentials (V_m_) were measured *in vitro* in thoracic flight muscle in a preparation that allowed for accurate control of the extracellular environment and temperature. Using this preparation, we tested the isolated and interactive effects of temperature (i.e. 0 and 20°C) and Na^+^/K^+^-ATPase inhibition (i.e. with or without ouabain) on muscle V_m_ in a complete cross-over experimental design for each of the ten *Drosophila* species (**Fig. 1**). Importantly, extracellular ion concentrations remained stable and within “normal” range in all experiments (see Materials and Methods).

**Figure 1.**
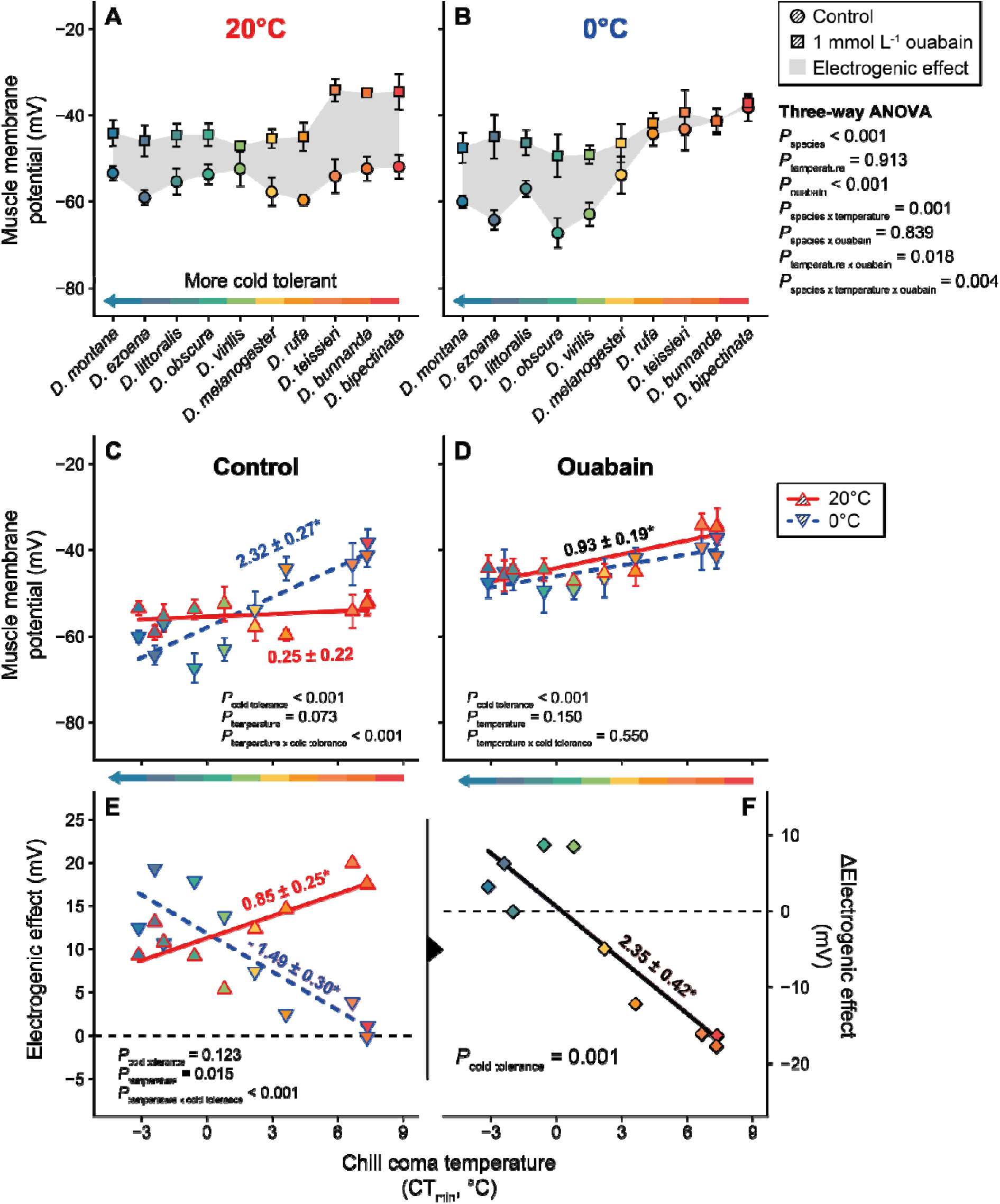
Cold tolerant *Drosophila* species are able to maintain muscle membrane potential and maintained a strong electrogenic effect at low temperature. **A** and **B**) Muscle V_m_ measured in thoracic flight muscle of 10 *Drosophila* species with different cold tolerances using an *in vitro* flight muscle preparation in the absence (circles) and presence (squares) of ouabain at permissive (**A**, 20°C) and low (**B**, 0°C) temperatures. The shaded grey area between the control- and ouabain treatments indicate the Na^+^/K^+^-ATPase-dependent electrogenic effect. **C**) Muscle V_m_ in the absence of ouabain at 20°C (full line and upwards-facing triangles in red) and 0°C (dashed line and downwards-facing triangles in blue) plotted against species cold tolerance (CT_min_ data from Kellermann et al. (2012)) showed a strong correlation between cold tolerance and muscle V_m_ at 0°C (slope = 2.32 ± 0.27, P < 0.001), while there was no relation at 20°C (slope = 0.25 ± 0.22, P = 0.271). **D**) In the presence of ouabain (at the same temperatures) there only was a main effect of cold tolerance which resulted in the positive correlation between muscle V_m_ and CT_min_ (i.e. a negative correlation between V_m_ and cold tolerance) (slope based on both temperatures = 0.93 ± 0.19, P < 0.001). **E**) The electrogenic effect was calculated as the difference between control and ouabain-treated preparations plotted against species CT_min_ showed a positive relation at 20°C (slope = 0.85 ± 0.25, P = 0.008) indicating that cold sensitive species have higher electrogenic effects at benign temperature. Conversely there was a significant negative association between electrogenic effect and CT_min_ at 0°C (slope = - 1.49 ± 0.30, P < 0.001) indicating that cold sensitive species have lower electrogenic effects at low temperature. These effects are evident from the significant main effect of species cold tolerant and the interaction with temperature. The dashed horizontal line indicates complete absence of an electrogenic effect. **F**) The change in the magnitude of the electrogenic effect between 20 and 0°C (ΔElectrogenic effect) plotted against species cold tolerance (CT_min_). The negative correlation between ΔElectrogenic effect and CT_min_ (slope = - 2.35 ± 0.42, P = 0.001) indicates that cold tolerant species (low CT_min_) are better at defending (or even potentiating) the electrogenic effect at low temperature. The dashed horizontal line indicates no change in electrogenic effect caused by the temperature decrease from 20 to 0°C. Points and error bars in panels A-D depict means ± s.e.m. based on 4-10 samples for each species/temperature/ouabain treatment, while points in E and F represent calculated means. *P* values depicted in C-F are from linear models and indicate the main and interactive effects of the factors denoted in subscript. Linear regression slopes are indicated next to their respective regression lines, and an asterisk (*) denotes a statistically significant correlation.

In our first analysis of this dataset (**Fig. 1A,B**), we found a significant effect of species (F_9,264_ = 9.4, P < 0.001) and a depolarizing effect of ouabain (F_1,264_ = 116.2, P < 0.001) but, surprisingly, no significant main effect of temperature (F_1,264_ = 0.0, P = 0.913). However, there was a significant three-way interaction between all factors (species x temperature x ouabain: F_9,264_ = 2.7, P = 0.004) as well as significant interactions between experimental temperature and species (F_9,264_ = 3.2, P = 0.001) and experimental temperature and ouabain (F_1,264_ = 5.7, P = 0.018) preventing accurate interpretations of the main effects. These interactions suggest that muscle V_m_ of different species is affected differently by temperature and that temperature influences how ouabain depolarizes V_m_ among species. Specifically, the data indicate that chill tolerant species were able to maintain a highly polarized muscle V_m_ at low temperature whereas the low temperature depolarized V_m_ in a number of chill sensitive species (i.e. compare circle symbols in **Fig. 1A** and **B**). Furthermore, the effect of ouabain at low temperature was greatly reduced in chill sensitive species but remained strong in a number of the more tolerant species (**Fig. 1B**).

To tease apart the effects of species, temperature and ouabain on muscle V_m_, we split the dataset into control and ouabain treatments and correlated these measurements to the species-specific chill tolerance (chill coma temperature, CT_min_). Doing so we found a strong significant effect of chill tolerance (F_1,147_ = 50.9, P < 0.001) on muscle V_m_ while the effect of the experimental temperature remained below the level of significance (F_1,147_ = 3.3, P = 0.073). Again, this pattern was confounded by a significant interaction between chill tolerance and experimental temperature (F_1,147_ = 35.1, P < 0.001) (**Fig. 1C**). Thus, at 20°C, a permissive temperature for all species, we found no significant relationship between muscle V_m_ and species chill tolerance (CT_min_) with an average V_m_ of approximately -56 mV. The linear regression of V_m_ against CT_min_ was not significantly different from 0 (slope = 0.25 ± 0.22 mV per °C CT_min_, P = 0.271, red line in **Fig. 1C**). In contrast, we found a strong and significant correlation between V_m_ and chill tolerance at 0°C (slope = 2.32 ± 0.27 mV per °C CT_min_, P < 0.001, blue dashed line in **Fig. 1C**). In summary, chill tolerant species were able to maintain muscle V_m_, or even polarize V_m_ further, at low temperatures while the sensitive species experienced severe depolarization.

When we examined the relationship of V_m_ against chill tolerance in the presence of ouabain (i.e. in the absence of the electrogenic effect generated by the Na^+^/K^+^-ATPase) (**Fig. 1D**) we found no effects of experimental temperature (temperature F_1,149_ = 2.1, P = 0.150) but a significant positive correlation between V_m_ and CT_min_ (slope = 0.93 ± 0.19 mV per °C CT_min_, P < 0.001, **Fig. 1D**). This shows that chill tolerant drosophilids have a more polarized muscle V_m_ in the absence of the electrogenic effect of the Na^+^/K^+^-ATPase regardless of the temperature. Importantly, in the presence of ouabain we found no interaction between experimental temperature and species chill tolerance (interaction F_1,149_ = 0.4, P = 0.550).

To further focus our analysis, we calculated the mean electrogenic effect for each species at the two temperatures and correlate these values to species-specific chill tolerance (**Fig. 1E**). Here we find a clear effect of temperature on the electrogenic effect (F_1,16_ = 7.4, P = 0.015), no effect of the CT_min_ (F_1,16_ = 2.6, P = 0.123), but again we also find a strong interaction between the two factors (F_1,16_ = 36.8, P < 0.001). Thus, this analysis confirms that exposure to low temperature strongly reduces the electrogenic effect in chill sensitive species, while chill tolerant species conversely experience a small increase in the electrogenic effect. To illustrate the interaction between temperature and the electrogenic effect we calculated the magnitude of the temperature-induced change in the electrogenic effect (i.e. the change in electrogenic effect between at 20 and 0°C) against species-specific chill tolerance (**Fig. 1F**) and demonstrate a strong positive correlation between species chill tolerance (i.e. a low CT_min_ implies high chill tolerance) and the ability to maintain or reinforce the electrogenic effect at low temperature (F_1,8_ = 31.3, P = 0.001).

### Metabolic rate estimates

To investigate if the superior maintenance of membrane potential and electrogenic potential at low temperature in chill tolerant *Drosophila* species came at a metabolic cost, we estimated the effects of temperature on the metabolic rate of all ten *Drosophila* species. This was done through measurements of CO_2_ production (V[CO_2_) at eight experimental temperatures ranging from 0 to 19°C. (**Fig. 2A**). The statistical analysis revealed main effects of species (F_9,415_ = 25.6, P < 0.001) and temperature (F_1,415_ = 14199.1, P < 0.001) but also a significant interaction between these factors (F_9,415_ = 26.8, P < 0.001), suggesting that the change in V[CO_2_ with temperature differs among species. As seen in **Fig. 2A** the mass adjusted V[CO_2_ ranged from ∼ 1.3 to ∼ 2.3 μL CO_2_ mg^−1^ h^−1^ at 19°C and decreased exponentially as a function of temperature to between 0.1 and 0.3 μL CO_2_ mg^−1^ h^−1^ at 0°C (note the log-transformed ordinate axis). Interestingly, there was a pattern to suggest that the chill sensitive species, that enter chill coma during the colder parts of the experiment experiments (*D. bipectinata*, *D. bunnanda*, *D. teissieri*, *D. rufa*, and *D. melanogaster*), had relatively higher V[CO_2_ at the low temperature. To partition the interactive effects of temperature and species we calculated Q_10_ values for each species and found this to range between 2.6 and 4.2 (**Fig 2B**, black symbols). When Q_10_ was correlated to species chill tolerance we found a significant negative relationship with species-specific chill coma temperatures (F_1,8_ = 18.6, P = 0.003; **Fig. 2B**) showing that more chill tolerant species had a larger reduction in V[CO_2_ in response to low temperatures (or *vice versa*). This trend was strengthened if Q_10_ values were calculated only over a low-temperature range (0 - 9°C) (F_1,8_ = 45.9, P < 0.001, blue line **Fig 2B**), and weakened below the level of statistical significance if calculated only for the higher temperatures (9 - 19°C)(F_1,8_ = 4.1, P = 0.077, red line **Fig 2B**). In accordance we found no support for the hypothesis that chill tolerant species spend excess energy at low temperature in order to defend their membrane potential and electrogenic effect. In fact, we observed, in contradiction to this hypothesis, that the species that defended or potentiated their electrogenic potential at low temperature, were also the species that experienced the largest decrease in metabolic rate at low temperature (see **Fig. S1**).

**Figure 2.**
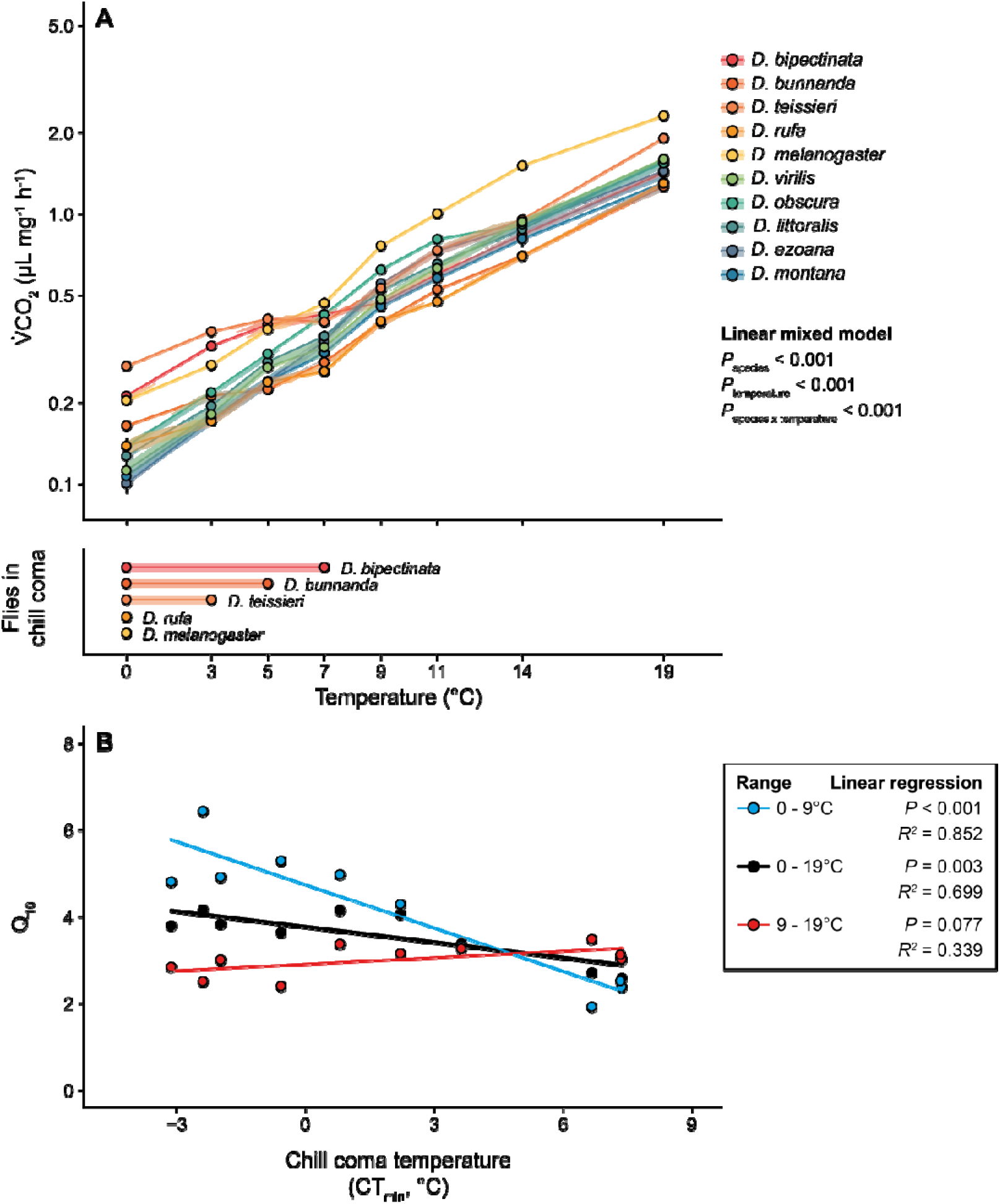
Increased thermal sensitivity of metabolism in cold tolerant *Drosophila* species. **A**) The effects of temperature on metabolism was estimated by measuring CO_2_ CO_2_) at temperatures from 0 to 19°C for ten *Drosophila* species with different cold tolerance, resulting in some species entering chill coma at lower temperatures. CO_2_ estimates. During this experiment, some species CO_2_ graph. **B**) Q_10_ values were calculated at low temperature where some species experience coma (≤ 9°C, blue), permissive temperatures (≥ 9°C, red), or over the entire temperature range of the experiment (0-19°C, black), and correlated to the species-specific thermal tolerance (here the chill coma temperature, from Kellermann et al. (2012)).

## Discussion

Progressive cooling challenges the capacity for homeostatic regulation of chill susceptible insects, leading to the depolarization of excitable tissues that causes a paralytic phenotype and ultimately also cell death, organismal injury, and mortality (Overgaard and MacMillan, 2017; Bayley et al., 2018; MacMillan, 2019; Overgaard et al., 2021; Robertson et al., 2023). Thus, defending membrane polarization during cold exposure is a key adaptation for chill tolerance in most insects and it is well known that cell depolarization occurs at different critical temperatures for different insect species (Hosler et al., 2000; Andersen et al., 2015; Andersen et al., 2017b). Cold-induced depolarization occurs due to a combination of a direct (thermodynamic) and indirect temperature effects including cold-induced changes in transmembrane ion balance (hyperkalemia) and reduced electrogenic polarization (Overgaard and MacMillan, 2017; Bayley et al., 2020; Bayley et al., 2021). Several studies have demonstrated how more tolerant species are better able to defend against loss of transmembrane ion balance during cold exposure (Overgaard and MacMillan, 2017; Overgaard et al., 2021), however, the variable role of the electrogenic component among species has so far received little attention.

In the present study we used a model system of ten *Drosophila* species with varying degrees of chill tolerance to investigate how the electrogenic potential generated by the Na^+^/K^+^-ATPase differed among species at both high (20°C) and low (0°C) temperature. The electrogenic contribution was evaluated through measurements of muscle membrane potential (V_m_) in the presence and absence of ouabain using an *in vitro* preparation exposed to “standard” *Drosophila* saline. Consequently, this approach allowed us to exclude any putative effects on V_m_ that could be related to the species’ different capacity to regulate hemolymph ion concentrations at low temperature.

The *in vitro* preparation used here is more invasive than *in vivo* preparations where the tissue is still surrounded by its own hemolymph. Nevertheless, we recorded muscle V_m_ of ∼ -55 mV at 20°C in standard saline for all 10 species (range -60 mV to -52 mV) (**Fig. 1A**) which resembles that previously found *in vivo* for five *Drosophila* species (-67 mV to -57 mV). This comparison includes two species (*D. montana* and *D. melanogaster*) represented by both *in vitro* (present study) and *in vivo* measurements (Andersen et al., 2015; Andersen and Overgaard, 2019) and therefore validates the experimental approach.

### Chill tolerant Drosophila maintain muscle V_m_ through preservation of the electrogenic effect

When examining the isolated effects of low temperature *in vitro* we found that chill tolerant species were better at protecting muscle V_m_ during cold exposure (Compare **Fig. 1A and Fig. 1B)** which matches our earlier *in vivo* observations for drosophilids (Andersen et al., 2015; Andersen and Overgaard, 2019), and other insects (Goller and Esch, 1990; Hosler et al., 2000; Andersen et al., 2017b). This observation underscores that the initial cold induced depolarization occurring in sensitive species is caused by factors unrelated to the changes in extracellular ion concentration (hyperkalemia) that are also hallmarks of chill sensitivity (Koštál et al., 2004; MacMillan and Sinclair, 2011b; MacMillan et al., 2014; Des Marteaux and Sinclair, 2016). Furthermore, this observation also highlights that tolerant species are able to mitigate this initial depolarization, which is, at least partially, linked to loss/preservation of an electric potential.

To explore directly the role of the Na^+^/K^+^-ATPase in the maintenance of muscle V_m_ through the electrogenic effect we performed experiment at high and low temperature with and without the addition of ouabain. At 20°C this resulted in a substantial depolarization in all species (**Fig. 1A**) (mean ± s.d. = 13.0 ± 4.5 mV, range = 5.3 – 20 mV). This confirms previous reports that insect muscle relies on a substantial electrogenic component of the muscle V_m_ (Rheuben, 1972; Wareham et al., 1974; Henon and Ikeda, 1981; MacMillan et al., 2014; Bayley et al., 2021). However, at low temperature ouabain had little to no effect in the chill sensitive species while the depolarizing effect remained in the chill tolerant species (**Fig. 1A-D**), resulting in a strong correlation between the Na^+^/K^+^-ATPase-dependent electrogenic effect and species-specific cold tolerance (**Fig. 1E,F**). Thus, we find strong support that chill tolerant *Drosophila* species maintain a considerable electrogenic polarization at low temperature while this is largely absent for the chill sensitive species. In other words chill sensitive species experience a temperature-induced loss of electrogenic polarization caused by reduced Na^+^/K^+^-ATPase-driven polarization.

### Why is the electrogenic effect maintained at low temperature in some species, but lost in others?

The electrogenic effect of the Na^+^/K^+^-ATPase stems from the net extrusion of a positive charge during each cycle (i.e. 3 Na^+^ out vs. 2 K^+^ in). The resulting current generates an electrical potential across the membrane that is proportional to both the magnitude of the current and the membrane resistance (*sensu* Ohm’s Law) (Thomas, 1972; Skou and Esmann, 1992). However, from our measurements it is not possible to determine if superior maintenance of electrogenic effect in cold tolerant species at low temperature is caused by relative high current (i.e. high Na^+^/K^+^-ATPase activity), high membrane resistance or through a combination. Below we briefly discuss possible factors that could contribute to the observed differences in electrogenic capacity among species.

The dipteran Na^+^/K^+^-ATPase is a heterodimer comprised of the catalytic α- and regulatory β-subunit (Emery et al., 1998); however, unlike mammals drosophilid flies have only one gene for the α-subunit (ATPα) and three isoforms of the β-subunit (nrv1-3), each of which have several transcript variations (Larkin et al., 2021), and the activity of the Na^+^/K^+^-ATPase is directly affected by the specific combination of subunits (Kawamura and Noguchi, 1991; Hilbers et al., 2016). It is possible that our results are linked to interspecific variation in the abundance of Na^+^/K^+^-ATPase subunits; however, to our knowledge this has not been investigated among *Drosophila* species. Two recent studies on thermal acclimation in *D. melanogaster* found no difference in abundance of the α-subunit between warm- and cold-acclimated conspecifics in brain and whole body samples, respectively (MacMillan et al., 2015c; Andersen et al., 2022). However, in the latter, an increased mRNA expression of a single β-subunit gene (nrv2) was found. How this particular change in subunit variation might affect transport rates and thermal sensitivity remains unknown.

The Na^+^/K^+^-ATPase activity is affected directly by the ion gradients against which it is operating (Glitsch, 2001) and by extension, it is highly dependent on available intracellular Na^+^ (and extracellular K^+^) for its activity (Everts and Clausen, 1992; Holmgren and Rakowski, 2006). Due to the finite reservoir of intracellular Na^+^, a continued electrogenic effect of the Na^+^/K^+^-ATPase is dependent on a continued electroneutral import of Na^+^ to fuel the pump. It was therefore suggested that membrane-bound Na^+^/K^+^/2Cl^−^ symporters were needed to support large electrogenic effects (Fraser and Huang, 2004; Bayley et al., 2021). While Na^+^/K^+^/2Cl^−^ symporters have not been investigated in relation to thermal tolerance and muscle V_m_, increased Na^+^/K^+^/2Cl^−^ symporter abundance has been demonstrated in the CNS of cold-acclimated locusts with improved neural cold tolerance and V_m_ maintenance (Srithiphaphirom et al., 2023).

Being a membrane-bound protein, the Na^+^/K^+^-ATPase is highly sensitive to changes in the membrane environment (Else and Wu, 1999), and numerous studies have documented membrane-lipid adaptations associated with differences in the thermal tolerance of insects (Koštál, 2010; Slotsbo et al., 2016). Currently, there is little-to-no direct evidence for a cause-effect relationship between membrane composition and V_m_ maintenance in insects. However, there are several reports of specific fatty acids and other membrane-integrating compounds modulating the activity of the Na^+^/K^+^-ATPase (Oishi et al., 1990; Jack-Hays et al., 1996). Future comparative and manipulative studies on the relation between membrane composition, membrane resistance, and Na^+^/K^+^-ATPase activity will likely provide valuable insights into the role of membrane viscosity in insect cold tolerance.

Separate from factors directly involved in the intrinsic transport rates of the Na^+^/K^+^-ATPase (i.e. the sections above) is its regulation by humoral factors (Ewart and Klip, 1995; Therien and Blostein, 2000). For example, the insects Na^+^/K^+^-ATPase has a number of non-catalytic phosphorylation sites, which can modulate its activity (Emery et al., 1998). Indeed, reversible phosphorylation lowers the activity of Na^+^/K^+^-ATPase in the goldenrod gall fly (*Eurosta solidaginis*) in a manner that matches how its activity is reduced during winter acclimatization and improvements in cold tolerance (McMullen and Storey, 2008). Again, regulatory signaling pathways have not been investigated in an interspecific model system of closely related species like the one used here, but we speculate that cold-tolerant species are able to maintain humoral stimulation at low temperature and therefore sustain Na^+^/K^+^-ATPase activity.

### Chill tolerant Drosophila species maintain muscle V_m_ independent of Na^+^/K^+^-ATPase electrogenesis at low temperature

In the absence of Na^+^/K^+^-ATPase-dependent electrogenesis the muscle V_m_ can be described by the diffusion potential, which largely depends on the transmembrane ion gradients and their relative permeabilities as described via the Goldman-Hodgkin-Katz equation (Goldman, 1943; Hodgkin and Katz, 1949). Under these conditions (i.e. in the presence of ouabain) we found that more cold tolerant *Drosophila* species were able to maintain a more polarized muscle V_m_ at both permissive (20°C) and low (0°C) temperature (**Fig. 1D**). The difference between the two temperatures can, if all else is equal, be described by the thermodynamic effect of temperature and should result in a ∼2.8 mV difference, which is in the same order of magnitude as what we find here (1.7 ± 1.7 mV, based on the linear model). Regardless, we kept the extracellular concentrations stable in our experimental preparation, and the differences observed between species here must relate to either differences in intracellular ion concentrations or their relative permeabilities. Cold acclimated *D. melanogaster* tend to have higher intracellular K^+^ concentrations than warm acclimated flies (Helou et al., 2024), which, if also present in more cold tolerant species, could partially drive the more polarized muscle V_m_ observed here. Additionally, a higher relative permeability for K^+^ has been indicated in locusts (Andersen et al., 2017a; Bayley et al., 2020), which would also tend to polarize muscle V_m_. If such mechanisms of intraspecific variation translate to our interspecific model system remains unknown, however, future studies could test this hypothesis.

### Maintained electrogenic effect at low temperature is not associated with increased energetic costs

Because the electrogenic effect is dependent on energetically demanding active ion transport, we hypothesized that preservation of electrogenic polarization in muscle of chill tolerant species came at a cost that was detectable in measurements of whole-animal CO_2_ production rate (V[CO_2_). Specifically, we expected that chill tolerant flies would have relatively higher metabolic rates at low temperatures compared to the chill sensitive species. Such a prediction follows from the popular, yet controversial, hypothesis of ‘metabolic cold adaptation’ (Fox, 1936; Addo□Bediako et al., 2002; Terblanche et al., 2009). Previous comparative studies of drosophila have not found any support for metabolic cold adaptation, but these previous studies have only investigated species differences at relatively high and non-stressful temperatures (above 10°C) where all species are able to survive and maintain homeostatic balance (Oikawa et al., 2006; Messamah et al., 2017).

Contrary to our hypothesis, and to the predictions of the ‘metabolic cold adaptation’ hypothesis, we find that chill-sensitive species from warmer climates tend to have relatively higher metabolic rates at the low temperature where they experience coma-inducing temperatures (**Fig. 2**). Thus sensitive species reduce metabolism less with decreasing temperature and have lower Q_10_ compared to the tolerant species that maintain polarization which experience reduction in metabolic rate with decreasing temperature (see **Figure S1**). This species difference in thermal sensitivity is particularly pronounced across the lower temperature range (blue points and line in **Fig. 2B**). From this we conclude that maintenance of the Na^+^/K^+^-ATPase-dependent electrogenic effect at low temperature is energetically cheap when evaluated at the whole organismal level. Conversely, we find that the inability to maintain electrogenic potential and functional neuromuscular performance in the chill sensitive species is not associated with a metabolic collapse. On the contrary these comatose flies spend relatively more energy in their cold stressed condition. Maintenance of metabolic rates and lowering of thermal sensitivities below the chill coma-inducing temperature has been reported previously for crickets (MacMillan et al., 2012), but the cause remains unexplored and unknown. The onset of chill coma is associated with a short burst of CO_2_ production which likely reflects the opening of spiracles and CO_2_ being released from the tracheal system (Stevens et al., 2010; Robertson et al., 2017); however, this momentary increase in CO_2_ release is unlikely to contribute to the elevated CO_2_ production observed over several hours in the small, comatose flies in our experiment.

## Conclusion

The principal finding of this study is the demonstration that the electrogenic effect of the Na^+^/K^+^-ATPase contributes an important and considerable component of the muscle V_m_ in all ten examined *Drosophila* species at 20°C. This electrogenic effect is highly sensitive to decreasing temperature in the most chill sensitive species, while tolerant species retain their electrogenic effect even at 0°C. Thus, the cold-induced muscle depolarization, that is a hallmark of chill susceptibility, is largely caused by loss of electrogenic polarization which, over time, is exacerbated by a subsequent disruption of ion balance (i.e. hyperkalemia) that further depolarizes the cells. The present study could not directly pinpoint why chill tolerant species are better able to maintain electrogenic polarization, but we speculate that this is linked to adaptations at the protein or membrane level. Regardless of the molecular underpinnings, we found no evidence for major metabolic costs in the species that maintain membrane polarization. In contrast we find cold-adapted species are able to maintain homeostatic balance and simultaneously have a lower metabolic turnover than their chill sensitive congeners.

## Data availability

All data has been made available to reviewers upon submission and will be linked with the article when published.

