## Supplementary materials for "Chill tolerant *Drosophila* species maintain electrogenic muscle membrane potential to resist cold-induced depolarization"

This document contains supplementary material for the manuscript titled:

by

Johannes Overgaard^1^, Jeppe Seamus Bayley^1^, Jacob Nørgaard Poulsen^1^, Nikolaj Johannes Skole Jensen^1^, Thomas Holm Pedersen^2^ and Jon Herskind^1^, Mads Kuhlmann Andersen^1,3^.

*^1^Department of Biology, Aarhus University, 8000 Aarhus C, Denmark*

*^2^Department of Biomedicine, Aarhus University, 8000 Aarhus C, Denmark*

*^3^Aarhus Institute for Advanced Studies, Aarhus University, 8000 Aarhus C, Denmark*

**Figure S1**

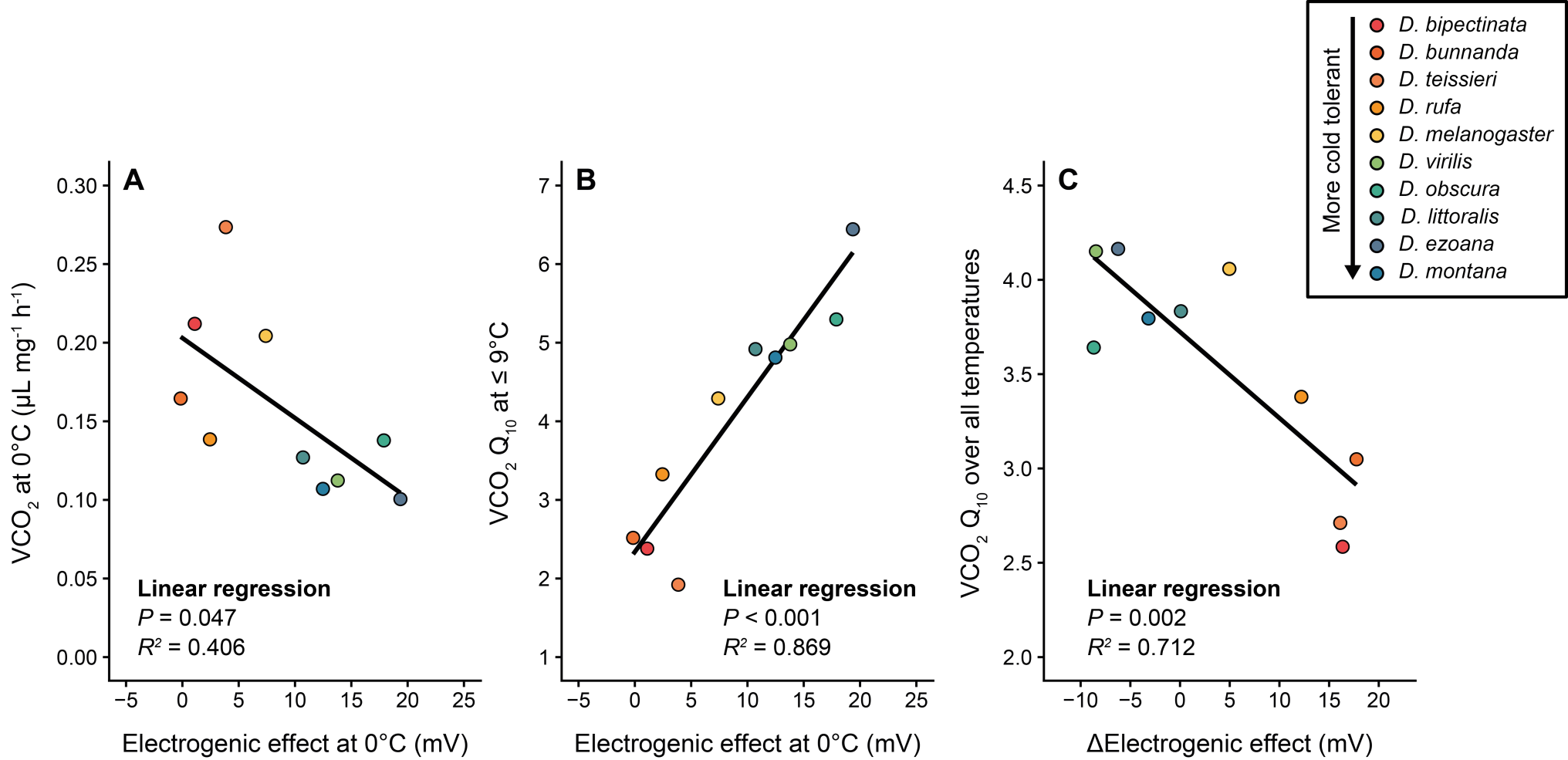

**Figure S1 – Correlations between the effects of temperature on species-specific CO_2_ production and maintenance of Na^+^/K^+^-ATPase-dependent electrogenic effect.** To test if the maintenance of muscle V_m_ at low temperature in cold tolerant *Drosophila* species came at a metabolic cost (i.e. the electrogenic Na^+^/K^+^-ATPase requires ATP to generate its current) we correlated **A**) the species-specific electrogenic effect at 0°C with CO_2_ production at 0°C, **B**) the species-specific electrogenic effect at 0°C with the Q_10_ of V̇CO_2_ at low temperatures (0 - 9°C), and **C**) the absolute reduction in electrogenic effects caused by the change in temperature from 20 to 0°C with the Q_10_ of V̇CO_2_ calculated over a similar temperature range (0 - 19°C). Points represent mean values and lines represent linear regression fits. Point colours indicate species chill susceptibility, with warmer colours indicating chill more sensitive species and colder colours indicating chill tolerant congeners (see the legend).

**Table S1**

This table contains information on the sample sizes for each species and temperature- and ouabain treatment from the membrane potential measurements, including the mean and standard error of the mean (depicted in the manuscript, Fig. 1A-D).

| **Species** | **Temperature [°C]** | **1 mmol L^-1^ ouabain [Y/N]** | **Sample size (N)** | **Mean muscle membrane potential (V_m_) [mV]** | **Standard error of the mean [mV]** |
| --- | --- | --- | --- | --- | --- |
| *Drosophila bipectinata* | 20 | N | 10 | -52,0 | 2,7 |
| *Drosophila bipectinata* | 20 | Y | 9 | -34,5 | 4,2 |
| *Drosophila bipectinata* | 0 | N | 8 | -38,2 | 3,1 |
| *Drosophila bipectinata* | 0 | Y | 8 | -37,1 | 1,7 |
| *Drosophila bunnanda* | 20 | N | 8 | -52,4 | 2,8 |
| *Drosophila bunnanda* | 20 | Y | 7 | -34,8 | 0,7 |
| *Drosophila bunnanda* | 0 | N | 8 | -41,2 | 2,7 |
| *Drosophila bunnanda* | 0 | Y | 8 | -41,3 | 2,9 |
| *Drosophila ezoana* | 20 | N | 8 | -59,1 | 1,7 |
| *Drosophila ezoana* | 20 | Y | 8 | -45,9 | 3,6 |
| *Drosophila ezoana* | 0 | N | 7 | -64,3 | 2,2 |
| *Drosophila ezoana* | 0 | Y | 8 | -45,0 | 5,1 |
| *Drosophila littoralis* | 20 | N | 8 | -55,4 | 2,9 |
| *Drosophila littoralis* | 20 | Y | 8 | -44,6 | 2,7 |
| *Drosophila littoralis* | 0 | N | 8 | -57,1 | 1,9 |
| *Drosophila littoralis* | 0 | Y | 8 | -46,4 | 2,9 |
| *Drosophila melanogaster* | 20 | N | 8 | -57,8 | 3,2 |
| *Drosophila melanogaster* | 20 | Y | 8 | -45,4 | 2,3 |
| *Drosophila melanogaster* | 0 | N | 7 | -53,9 | 4,3 |
| *Drosophila melanogaster* | 0 | Y | 8 | -46,5 | 4,5 |
| *Drosophila montana* | 20 | N | 7 | -53,5 | 1,7 |
| *Drosophila montana* | 20 | Y | 8 | -44,1 | 3,0 |
| *Drosophila montana* | 0 | N | 7 | -60,0 | 1,4 |
| *Drosophila montana* | 0 | Y | 8 | -47,5 | 3,5 |
| *Drosophila obscura* | 20 | N | 8 | -53,7 | 2,3 |
| *Drosophila obscura* | 20 | Y | 8 | -44,5 | 2,6 |
| *Drosophila obscura* | 0 | N | 8 | -67,3 | 3,4 |
| *Drosophila obscura* | 0 | Y | 6 | -49,4 | 5,0 |
| *Drosophila rufa* | 20 | N | 7 | -59,7 | 1,2 |
| *Drosophila rufa* | 20 | Y | 10 | -45,0 | 3,3 |
| *Drosophila rufa* | 0 | N | 8 | -44,3 | 2,8 |
| *Drosophila rufa* | 0 | Y | 8 | -41,8 | 2,4 |
| *Drosophila teissieri* | 20 | N | 8 | -54,2 | 3,9 |
| *Drosophila teissieri* | 20 | Y | 8 | -34,2 | 2,6 |
| *Drosophila teissieri* | 0 | N | 8 | -43,2 | 4,9 |
| *Drosophila teissieri* | 0 | Y | 7 | -39,4 | 5,2 |
| *Drosophila virilis* | 20 | N | 6 | -52,5 | 4,0 |
| *Drosophila virilis* | 20 | Y | 5 | -47,1 | 1,0 |
| *Drosophila virilis* | 0 | N | 4 | -63,0 | 2,7 |
| *Drosophila virilis* | 0 | Y | 5 | -49,2 | 2,2 |

**Table S2**

This table contains information on the sample sizes for each species and experimental temperature from the respirometry experiment, including the mean and standard error of the mean (depicted in the manuscript, Fig. 2A).

| **Species** | **Temperature [°C]** | **Sample size (N)** | **Mean mass adjusted, mass-specific CO2 production [μL CO_2_/mg/h]** | **Standard error of the mean [μL CO_2_/mg/h]** |
| --- | --- | --- | --- | --- |
| *Drosophila bipectinata* | 19 | 6 | 1,423 | 0,033 |
| *Drosophila bipectinata* | 14 | 6 | 0,839 | 0,022 |
| *Drosophila bipectinata* | 11 | 6 | 0,600 | 0,021 |
| *Drosophila bipectinata* | 9 | 6 | 0,469 | 0,016 |
| *Drosophila bipectinata* | 7 | 6 | 0,426 | 0,012 |
| *Drosophila bipectinata* | 5 | 6 | 0,390 | 0,019 |
| *Drosophila bipectinata* | 3 | 6 | 0,325 | 0,011 |
| *Drosophila bipectinata* | 0 | 6 | 0,212 | 0,009 |
| *Drosophila bunnanda* | 19 | 6 | 1,266 | 0,054 |
| *Drosophila bunnanda* | 14 | 6 | 0,702 | 0,027 |
| *Drosophila bunnanda* | 11 | 6 | 0,526 | 0,027 |
| *Drosophila bunnanda* | 9 | 6 | 0,397 | 0,015 |
| *Drosophila bunnanda* | 7 | 6 | 0,283 | 0,008 |
| *Drosophila bunnanda* | 5 | 6 | 0,225 | 0,005 |
| *Drosophila bunnanda* | 3 | 6 | 0,212 | 0,007 |
| *Drosophila bunnanda* | 0 | 6 | 0,165 | 0,007 |
| *Drosophila ezoana* | 19 | 7 | 1,445 | 0,089 |
| *Drosophila ezoana* | 14 | 4 | 0,909 | 0,054 |
| *Drosophila ezoana* | 11 | 6 | 0,731 | 0,023 |
| *Drosophila ezoana* | 9 | 6 | 0,554 | 0,023 |
| *Drosophila ezoana* | 7 | 6 | 0,338 | 0,015 |
| *Drosophila ezoana* | 5 | 6 | 0,241 | 0,011 |
| *Drosophila ezoana* | 3 | 6 | 0,173 | 0,008 |
| *Drosophila ezoana* | 0 | 6 | 0,101 | 0,008 |
| *Drosophila littoralis* | 19 | 6 | 1,579 | 0,031 |
| *Drosophila littoralis* | 14 | 12 | 0,880 | 0,026 |
| *Drosophila littoralis* | 11 | 6 | 0,655 | 0,030 |
| *Drosophila littoralis* | 9 | 12 | 0,522 | 0,020 |
| *Drosophila littoralis* | 7 | 6 | 0,353 | 0,019 |
| *Drosophila littoralis* | 5 | 6 | 0,283 | 0,018 |
| *Drosophila littoralis* | 3 | 6 | 0,195 | 0,017 |
| *Drosophila littoralis* | 0 | 6 | 0,127 | 0,016 |
| *Drosophila melanogaster* | 19 | 6 | 2,318 | 0,045 |
| *Drosophila melanogaster* | 14 | 12 | 1,515 | 0,021 |
| *Drosophila melanogaster* | 11 | 6 | 1,007 | 0,019 |
| *Drosophila melanogaster* | 9 | 12 | 0,764 | 0,020 |
| *Drosophila melanogaster* | 7 | 12 | 0,468 | 0,025 |
| *Drosophila melanogaster* | 5 | 6 | 0,374 | 0,017 |
| *Drosophila melanogaster* | 3 | 6 | 0,276 | 0,013 |
| *Drosophila melanogaster* | 0 | 6 | 0,204 | 0,008 |
| *Drosophila montana* | 19 | 5 | 1,308 | 0,084 |
| *Drosophila montana* | 14 | 5 | 0,811 | 0,049 |
| *Drosophila montana* | 11 | 5 | 0,579 | 0,017 |
| *Drosophila montana* | 9 | 5 | 0,455 | 0,021 |
| *Drosophila montana* | 7 | 5 | 0,306 | 0,015 |
| *Drosophila montana* | 5 | 5 | 0,239 | 0,013 |
| *Drosophila montana* | 3 | 5 | 0,176 | 0,009 |
| *Drosophila montana* | 0 | 5 | 0,107 | 0,006 |
| *Drosophila obscura* | 19 | 6 | 1,546 | 0,056 |
| *Drosophila obscura* | 14 | 4 | 0,919 | 0,038 |
| *Drosophila obscura* | 11 | 6 | 0,808 | 0,039 |
| *Drosophila obscura* | 9 | 6 | 0,624 | 0,028 |
| *Drosophila obscura* | 7 | 6 | 0,424 | 0,013 |
| *Drosophila obscura* | 5 | 6 | 0,304 | 0,010 |
| *Drosophila obscura* | 3 | 6 | 0,219 | 0,007 |
| *Drosophila obscura* | 0 | 6 | 0,138 | 0,005 |
| *Drosophila rufa* | 19 | 6 | 1,300 | 0,021 |
| *Drosophila rufa* | 14 | 12 | 0,698 | 0,012 |
| *Drosophila rufa* | 11 | 6 | 0,474 | 0,015 |
| *Drosophila rufa* | 9 | 12 | 0,401 | 0,011 |
| *Drosophila rufa* | 7 | 6 | 0,261 | 0,013 |
| *Drosophila rufa* | 5 | 6 | 0,239 | 0,007 |
| *Drosophila rufa* | 3 | 6 | 0,171 | 0,007 |
| *Drosophila rufa* | 0 | 6 | 0,139 | 0,009 |
| *Drosophila teissieri* | 19 | 5 | 1,913 | 0,067 |
| *Drosophila teissieri* | 14 | 4 | 0,955 | 0,044 |
| *Drosophila teissieri* | 11 | 6 | 0,735 | 0,026 |
| *Drosophila teissieri* | 9 | 6 | 0,532 | 0,020 |
| *Drosophila teissieri* | 7 | 6 | 0,399 | 0,008 |
| *Drosophila teissieri* | 5 | 6 | 0,409 | 0,021 |
| *Drosophila teissieri* | 3 | 6 | 0,367 | 0,018 |
| *Drosophila teissieri* | 0 | 6 | 0,274 | 0,011 |
| *Drosophila virilis* | 19 | 6 | 1,600 | 0,034 |
| *Drosophila virilis* | 14 | 11 | 0,937 | 0,025 |
| *Drosophila virilis* | 11 | 5 | 0,631 | 0,027 |
| *Drosophila virilis* | 9 | 11 | 0,483 | 0,015 |
| *Drosophila virilis* | 7 | 5 | 0,321 | 0,024 |
| *Drosophila virilis* | 5 | 6 | 0,271 | 0,013 |
| *Drosophila virilis* | 3 | 6 | 0,181 | 0,011 |
| *Drosophila virilis* | 0 | 6 | 0,112 | 0,007 |
